# Genome-informed trophic classification and functional characterization of virulence proteins from the maize tar spot pathogen *Phyllachora maydis*

**DOI:** 10.1101/2024.01.22.576543

**Authors:** Abigail Rogers, Namrata Jaiswal, Emily Roggenkamp, Hye-Seon Kim, Joshua S. MacCready, Martin I. Chilvers, Steven R. Scofield, Anjali S. Iyer-Pascuzzi, Matthew Helm

**Affiliations:** Department of Botany and Plant Pathology, Purdue University West Lafayette, IN 47907, U.S.A; Crop Production and Pest Control Research Unit, U.S. Department of Agriculture-Agricultural Research Service (USDA-ARS), West Lafayette, IN 47907, U.S.A; Department of Plant, Soil, and Microbial Sciences, Michigan State University, East Lansing, MI, 48824, U.S.A; U.S. Department of Agriculture-Agricultural Research Service (USDA-ARS), National Center for Agricultural Utilization Research, Mycotoxin Prevention and Applied Microbiology Research Unit, Peoria, IL 61604, U.S.A

**Keywords:** *Phyllachora maydis*, tar spot, CAZymes, effectors, effector localization, immune suppression, reactive oxygen species

## Abstract

*Phyllachora maydis* is an ascomycete foliar fungal pathogen and the causal agent of tar spot in maize. Though *P. maydis* is considered one of the most economically important foliar pathogens of maize, our general knowledge of the trophic lifestyle and functional role of effector proteins from this fungal pathogen remains limited. Here, we utilized a genome-informed approach to predict the trophic lifestyle of *P. maydis* and functionally characterized a subset of candidate effectors from this fungal pathogen. Leveraging the most recent *P. maydis* genome annotation and the CATAStrophy pipeline, we show this fungal pathogen encodes a predicted Carbohydrate-active enzymes (CAZymes) repertoire consistent with that of biotrophs (monomertrophs). To investigate fungal pathogenicity, we selected eighteen candidate effector proteins that were previously shown to be expressed during primary disease development. We assessed whether these putative effectors share predicted structural similarity with other characterized fungal effectors and determined whether any suppress plant immune responses. Using AlphaFold2 and Foldseek, we showed one candidate effector, PM02_g1115, adopts a predicted protein structure similar to that of an effector from *Verticillium dahlia*. Furthermore, transient expression of candidate effector-fluorescent protein fusions in *Nicotiana benthamiana* revealed that most effector proteins localize to both the nucleus and the cytosol. Importantly, three candidate effectors consistently attenuated chitin-mediated reactive oxygen species production in *N. benthamiana*. Collectively, these results presented herein provide valuable insights into the predicted trophic lifestyle and putative functions of effectors from *P. maydis* and will likely stimulate continued research to elucidate the molecular mechanisms used by *P. maydis* to induce tar spot.

## INTRODUCTION

Plants have evolved a two-tiered innate immune system that is activated, in part, upon the recognition of pathogen-associated molecular patterns (PAMPs) by cell surface-localized pattern-recognition receptors (PRRs) (Chen et al., 2022; McCombe et al., 2022; Rhodes et al., 2022). Upon the perception of PAMPs at the cell surface, plant PRRs initiate a coordinated defense response referred to as pattern-triggered immunity (PTI) (Chen et al., 2022; McCombe et al., 2022; Rhodes et al., 2022). PTI is partially characterized by an increase in reactive oxygen species (ROS) production, activation of mitogen-activated protein kinase (MAPK) signaling cascades, calcium influx and accumulation within the host cytosol, and stomatal closure (Chen et al., 2022; Jones and Dangl, 2006; McCombe et al., 2022; Rhodes et al., 2022). To circumvent and evade these cell surface-triggered immune responses, pathogens secrete virulence molecules, often proteins, known as effectors to either the host apoplast or cytosol to evade the host-triggered immune responses (Bentham et al., 2020; Jones and Dangl, 2006; Ngou et al., 2022). For example, the apoplastic effector Pep1 from the maize smut fungus *Ustilago maydis* functions as a peroxide inhibitor, thereby suppressing extracellular ROS production and promoting disease progression (Hemetsberger et al., 2012). Importantly, loss of Pep1 protein expression leads to a significant reduction in the virulence of *U. maydis* as Pep1 mutants cannot penetrate host cells to advance the infection cycle (Doehlemann et al., 2009). Likewise, Saado and colleagues (2022) recently showed the *U. maydis* effector Rip1 interferes with host ROS burst and such immune suppression activity was independent of its *in planta* subcellular localization. In addition, two effector proteins from the Southern corn rust pathogen, *Puccinia polysora*, were recently identified and functionally characterized (Chen et al., 2022; Deng et al., 2022). Recent work from Chen et al., (2022) showed that the *P. polysora* AvrRppK effector suppresses multiple pattern-triggered immune responses including MAPK activity and ROS production. Importantly, transgenic maize constitutively expressing AvrRppK showed enhanced susceptibility to *P. polysora* demonstrating this fungal effector attenuates host immune response and likely has a functional role in Southern corn rust disease development (Chen et al., 2022). Additionally, Deng and colleagues (2022) identified 120 predicted secreted effectors from an avirulent *P. polysora* strain. Infiltration of purified protein from one candidate effector protein, AvrRppC, into a *P. polysora*-resistant maize inbred resulted in activation of robust immune responses in maize (Deng et al., 2022).

In addition to secreting effectors, fungal pathogens often utilize Carbohydrate-active enzymes (CAZymes) to further facilitate disease development (Bradley et al., 2022; Zhao et al., 2013). CAZymes are a broad class of enzymes that often have a functional role in the biosynthesis or degradation of complex carbohydrates, including plant cell wall degradation (Bradley et al., 2022; Zhao et al., 2013). CAZymes can be classified into six superfamilies: auxiliary activity (AA) enzymes, carbohydrate-binding modules (CBMs), glycoside hydrolases (GHs), glycosyl transferases (GTs), carbohydrate esterases (CEs), and polysaccharide lyases (PLs) (Bradley et al., 2022; Zhao et al., 2013). Furthermore, several studies have reported the fungal secretome composition, including the predicted secreted CAZyme repertoire, is often strongly associated with trophic lifestyle (Hane et al., 2020; Jia et al., 2023; Lowe and Howlett, 2012). For example, comparative genome analyses revealed hemibiotrophic fungal pathogens encode, on average, a total of 259 predicted secreted CAZymes (Jia et al., 2023). Necrotrophic fungi usually contain a total of 200 predicted secreted CAZymes, while biotrophs encode 65 predicted secreted CAZymes (Jia et al., 2023). Additionally, biotrophic fungi and oomycetes often contain ∼100 GH-encoding genes while hemibiotrophic and necrotrophic fungi have ∼300 GH-encoding genes (Hane et al., 2020; Wang et al., 2023; Zerillo et al., 2013; Zhao et al., 2013). Hence, the quantity and type of CAZyme-encoding genes can often assist in inferring the lifestyle and trophic classification of a fungal pathogen. Using these observations, Hane et al., (2020) developed a bioinformatic pipeline to predict the lifestyle and trophic phenotype of novel filamentous fungal pathogens. This method, referred to as CAZyme-Assisted Training And Sorting of-trophy (CATAStrophy), predicts the lifestyle (saprotroph, monomertroph [corresponding to biotrophs], mesotroph [corresponding to hemibiotrophs], vasculartroph, and polymertroph [corresponding to necrotrophs]) of filamentous fungal pathogens exclusively based on the fungal CAZyme composition (Hane et al., 2020).

*Phyllachora maydis* is an ascomycete foliar pathogen of maize (*Zea mays* sp. *mays*) that causes tar spot disease (Ruhl et al., 2016; Rocco da Silva et al., 2021; Valle-Torres et al., 2020). Since its discovery in the United States in 2015, *P. maydis* has spread significantly and most commercial maize hybrids are susceptible to this fungal pathogen (Mueller et al., 2022; Rocco da Silva et al., 2021; Valle-Torres et al., 2020).

Moreover, elucidating the trophic lifestyle as well as the molecular mechanisms that contribute to *P. maydis* pathogenicity remains challenging as its reported obligate lifestyle hinders genetic manipulation (Helm et al., 2022; Rocco da Silva et al., 2021; Telenko et al., 2020). MacCready et al., (2023) recently reported a high-quality *P. maydis* isolate PM02 genome (assembly ASM2933922v1) and accompanying transcriptome from primary tar spot lesions during early disease development. The genome size of *P. maydis* genome is ∼64 Mbp and encodes 9,630 predicted proteins, of which 492 proteins are predicted to be secreted (MacCready et al., 2023). Within the *P. maydis* secretome, 163 proteins are predicted to encode effector-like sequences (MacCready et al., 2023). Intriguingly, six of the top twenty-five most abundantly expressed genes are putative effectors, suggesting this fungal pathogen utilizes effectors to attenuate immune responses during primary disease development (MacCready et al., 2023). Recent work by Caldwell and colleagues (2023) has also provided significant insights into the infection strategy used by *P. maydis* to colonize maize foliar leaf tissue. Histological-based, high-resolution light and electron microscopy analyses revealed that *P. maydis* hyphae penetrate the host epidermis and subsequently spread intracellularly and intercellularly (Caldwell et al., 2023). Once established, this fungal pathogen forms four distinct sub-structures within the host tissue: vegetative hyphae, clypeus, pycnidium, and perithecia (Caldwell et al., 2023). Hence, these studies reveal that *P. maydis* likely uses effector proteins to help establish and facilitate infection within host tissues (Caldwell et al., 2023; MacCready et al., 2023).

Here, we utilized a genome-informed approach to predict the trophic lifestyle of *P. maydis* and functionally characterized a subset of candidate effectors from *P. maydis*. We screened the predicted secretome of *P. maydis* using the CAZyme-Assisted Training And Sorting of-trophy (CATAStrophy) pipeline and identified 92 predicted secreted CAZymes, of which 57 are annotated as glycoside hydrolases. Based on the quantity and type of predicted secreted CAZymes, we concluded that *P. maydis* is likely a monomertroph (biotroph). To further investigate fungal pathogenicity, we selected eighteen candidate effector proteins, which have been previously shown to be expressed during disease development (MacCready et al., 2023). Specifically, we assessed whether any these putative effectors share structural similarity with other characterized fungal effectors, determined the *in planta* subcellular localizations, and tested whether any candidate effectors attenuate chitin-mediated ROS production in *N. benthamiana*. Using advanced protein structure prediction algorithms, we show that one candidate effector, PM02_g1115, adopted a predicted protein structure similar to that of the PevD1 effector from *Verticillium dahlia*. Using *Agrobacterium tumefaciens*-mediated heterologous expression in *Nicotiana benthamiana*, we show that most of the candidate effector proteins localize to both the nucleus and the cytosol. However, fluorescence signal from two candidate effectors, PM02_g378 and PM02_g2610, accumulated predominantly in the cytosol with weak fluorescence detected in the nucleus. Importantly, three candidate effectors consistently attenuated chitin-mediated reactive oxygen species production. Collectively, our results suggest *P. maydis* encodes CAZymes consistent with that of monomertrophs and utilizes candidate effector proteins to suppress immune responses. The results provided here will likely stimulate new research aimed at elucidating the host biological processes manipulated by *P. maydis* to induce disease.

## MATERIALS AND METHODS

### Prediction of *P. maydis* CAZymes and trophic classification

The CATAStrophy software v.0.1.0 was used to predict the carbohydrate active enzymes (CAZyme) repertoire and infer the trophic lifestyle of *Phyllachora maydis* (Hane et al., 2020). Briefly, we leveraged the PM02 genome annotation from NCBI genome assembly ASM2933922v1 and executed the CATAStrophy pipeline, which uses the dbCAN CAZyme (v10) database and HMMER3 (v.3.3.2) to predict the secreted CAZyme repertoire (MacCready et al., 2023; Zheng et al., 2023; hmmer.org). The –counts option was used to approximate the number of each group of CAZyme annotation. The trophic lifestyle was inferred by the quantity of the different types of predicted CAZymes and reported as a relative centroid distance (RCD) value, which is the relative proportion between 0 (farthest) and 1 (closest) to show the likelihood of a fungal pathogen being within a specific trophic classification (Hane et al., 2020).

### Selection of candidate effectors from *P. maydis* isolate PM02 (assembly ASM2933922v1)

Candidate effectors were selected by mining an improved genome assembly of *P. maydis* (assembly ASM2933922v1) (isolate PM02)and accompanying transcriptome (BioProject PRJNA928553) for proteins that encoded effector-like sequence characteristics, and which are expressed during primary disease formation (Helm et al., 2022; MacCready et al., 2023). Overall, we selected the top eighteen genes that fulfilled our selection criteria (Table 2; Supplemental Table 1). Signal peptides and effector predictions were verified using SignalP (v6.0) (https://services.healthtech.dtu.dk/service.php?SignalP) and EffectorP (v3.0) (http://effectorp.csiro.au/), respectively (Sperschneider and Dodds, 2022; Teufel et al., 2022). LOCALIZER (v1.0) ((http://localizer.csiro.au/) was used to identify potential nuclear localization signals (NLS) (Sperschneider et al., 2017). Putative protein-encoding domains were identified with the InterPro Database (https://www.ebi.ac.uk/interpro/) using the candidate effector open reading frame as a query.

### Prediction of *P. maydis* effector protein structures using AlphaFold2

The protein structures of mature *P. maydis* candidate effectors (without their signal peptide sequences) were predicted using the ColabFold v1.5.2: AlphaFold2 using MMseqs2 with default parameters (Mirdita et al., 2022). The ColabFold Jupyter Notebook and codes (https://github.com/sokrypton/ColabFold/blob/main/beta/AlphaFold2_advanced.ipynb) were used to visualize the predicted tertiary structures of each effector protein with the confidence prediction .pdb files and the model .pkl files (Mirdita et al., 2022). Predicted protein structures with the highest Predicted Local Distance Difference Test (pLDDT) confidence scores were captured and visualized using ChimeraX-1.5 (Meng et al., 2023). To identify proteins with putative structural similarity to the selected candidate effectors, we leveraged the Foldseek webserver (https://search.foldseek.com/search). The PDB file with the highest confidence prediction as determined by AlphaFold2 (i.e. ranked_0.pdb) was used as the query. Search parameters for the Foldseek search were set such that all publicly available structure databases (AlphaFold/Uniprot50-v4, AlphaFold/Swiss-Prot-v4, AlphaFold/Proteome-v4, CATH50-v4.3.0, MGnify-ESM30-v1, PDB100, and GMGCL-2204) were selected and the taxonomic filer was set to “Fungi” (Mode: 3Di/AA) (Barrio-Hernandez et al., 2023; van Kempen et al., 2023).

### Plant growth conditions

Seeds of *Nicotiana benthamiana* were sown in plastic pots containing Berger Seed and Propagation Mix supplemented with Osmocote slow-release fertilizer (14-14- 14) and maintained either in a growth chamber with a 16:8 h photoperiod (light:dark) at 24°C with light and 20°C in the dark, with average light intensities at plant height of 120 µmols/m^2^/s, or grown at room temperature with light intensity between 120 −140 μmols/m^2^/s (Philips F32T8/L941 Deluxe Cool White bulbs) and a 16:8 h photoperiod.

### Generation of plant expression constructs

All plant expression constructs were generated using a method previously described by Helm et al., (2022). Briefly, a commercial gene synthesis provider was used to synthesize the open reading frame (ORF) encoding the mature effector (i.e. lacking the predicted signal peptide). The *att*L1 and *att*L4 Gateway sequences were added to the 5’ and 3’ ends, respectively, of the ORF and subsequently inserted into pUC57 by the service provider. The resulting plasmids, pBSDONR(L1-L4):PmECs, were mixed with pBSDONR(L4r-L2):sYFP (Helm et al., 2022; Qi et al., 2012) and recombined into the Gateway destination vector pEarleyGate100 (pEG100) (Earley et al., 2006; Helm et al., 2022) using LR Clonase II (Invitrogen), thereby generating C-terminal-tagged effector-fluorescent protein fusions.

### Transient protein expression in *Nicotiana benthamiana*

Transient expression assays were performed as previously reported by Helm et al., (2022) with minor modifications. The candidate effector-fluorescent protein fusions were mobilized into *Agrobacterium tumefaciens* GV3101 (pMP90) and grown overnight in LB media supplemented with 25µg of gentamicin sulfate and 50µg of kanamycin per milliliter at 30°C. Following overnight incubation, cells were pelleted by centrifuging at 3200rpm for 3 minutes at room temperature. The bacterial pellet was resuspended in 10mM MgCl_2_, adjusted to an optical density at 600nm (OD_600_) between 0.5 and 0.6, and incubated with 100µM of acetosyringone for 2-4 hours at room temperature. Bacterial suspensions were infiltrated into the abaxial side of *N. benthamiana* leaves using a needless syringe.

### Confocal microscopy

Confocal microscopy of *N. benthamiana* epidermal cells was performed twenty-four hours post-agroinfiltration using a Zeiss LSM880 Axio Examiner upright confocal microscope with a Plan Apochromat 20x/0.8 objective, as previously described (Helm et al., 2022). Image analysis was performed using Zeiss Zen Blue Lite.

### Immunoblot analyses

Agroinfiltrated *N. benthamiana* leaf tissue was collected, and flash frozen in liquid nitrogen twenty-four and forty-eight hours post-agroinfiltration, and total protein was extracted as previously described by Helm et al., (2022) for immunoblot analyses. Ten microliters of isolated total protein were separated on a 4-20% Tris-glycine stain free polyacrylamide gel (Bio-Rad) at 180 V for one hour in 1X Tris/glycine/SDS running buffer. Total proteins were subsequently transferred to nitrocellulose membranes (GE Water and Process Technologies) and stained with PonceauS to confirm transfer and equal loading of proteins. Membranes were washed with 1X Tris-buffered saline (50mM Tris-HCl, 150mM NaCl, pH 7.5) solution supplemented with 0.1% Tween20 (TBST) and blocked in 5% skim milk (Becton, Dickinson & Company) for at least 1 hour at room temperature. Proteins were detected with horseradish peroxidase (HRP)-conjugated anti-GFP antibody (1:5,000; Miltenyi Biotec) for one hour at room temperature. Membranes were washed in 1X TBST solution and subsequently incubated with Clarity Western ECL (Bio-Rad) substrate solution for 5 minutes. Imaging of immunoblots was performed with an ImageQuant 500 CCD imaging system.

### Luminol-based ROS assay in *N. benthamiana*

Attenuation of ROS production in *N. benthamiana* was performed as previously described using a luminol-based chemiluminescence assay and chitin as an elicitor (Jaiswal et al., 2022). *N. benthamiana* leaf discs (5mm diameter) transiently expressing either free sYFP (3xHA:sYFP; Helm et al., 2022) or the candidate effector-fluorescent protein fusions were sampled two days post-agroinfiltration using a cork borer, washed three times in deionized water, and incubated overnight in sterile water in a 96-well OptiPlate^TM^ microplate (Perkin Elmer). The following day, the deionized water was replaced with chitin elicitation solution (luminol [30μg/mL], horseradish peroxidase [20μg/mL], chitin [hexamer] [5μg/mL], and nuclease-free water) and ROS production was monitored by chemiluminescence for 40 minutes in a microplate reader (Tecan Infinite M200 Pro), with higher luminescence values indicating greater ROS production. Experiments were performed three independent times.

## RESULTS

### Prediction of the CAZyme repertoire and trophic classification of *P. maydis*

The type and quantity of Carbohydrate-active enzymes (CAZymes) encoded by fungal phytopathogens is often correlated with lifestyle and infection strategy (Hane et al., 2020). Hence, analysis of CAZyme-encoding genes can assist in inferring the lifestyle and trophic classification of fungal pathogens. Current knowledge, albeit anecdotally, suggests *P. maydis* is an obligate biotroph as it likely requires photosynthetically active leaf tissue for growth and reproduction, and efforts to culture this fungal pathogen thus far have been unsuccessful (Rocco da Silva et al., 2021; Solórzano et al., 2023). We, therefore, screened the *P. maydis* proteome using the CAZyme-Assisted Training And Sorting of-trophy (CATAStrophy) prediction tool to predict the trophic lifestyle of *P. maydis* exclusively based on the composition of CAZyme-encoding genes (Hane et al., 2020). These *in silico* analyses revealed that this filamentous fungal pathogen encodes 336 predicted CAZymes, of which 92 encode signal peptides and are thus putatively secreted (Table 1A). Upon further examination of the *P. maydis* CAZyme repertoire, we found this fungal pathogen contains 54 auxiliary activity enzymes, 20 carbohydrate-binding modules, 153 glycoside hydrolases, 78 glycosyl transferases, 31 carbohydrate esterases, and no polysaccharide lyases (Table 1A). Furthermore, of the 92 predicted secreted CAZymes, 18 are annotated as auxiliary activity enzymes, 7 as carbohydrate-binding modules, 57 as glycoside hydrolases, 3 as glycosyl transferases, and 7 as carbohydrate esterases. Importantly, the CATAStrophy platform classified *P. maydis* as having a trophic lifestyle consistent with that of non-haustorial monomertrophs (biotrophs) based on the quantity and type of predicted secreted CAZyme-encoding genes (Table 1B). The relative centroid distance (RCD) values for saprotroph, mesotroph, polymertroph, and vasculartroph were 0.88, 0.42, 0.43, and 0 respectively (Table 1B).

**Table 1.** CAZyme annotation and predicted trophic classification of *P. maydis* isolate PM02 (assembly ASM2933922v1) as determined by CATAStrophy. A) *P. maydis* encodes 336 predicted CAZymes, of which 92 are putatively secreted CAZymes. The CATAStrophy (v.0.1.0) software was used to predict the carbohydrate active enzymes (CAZyme) repertoire. AA, Auxiliary activity enzymes; CBMs, carbohydrate-binding modules; GHs, glycoside hydrolases; GTs, glycosyl transferases; CEs, carbohydrate esterases; PLs, polysaccharide lyases. **B)** Summary of trophic classification predictions for *P. maydis* as determined by CATAStrophy. The predicted secreted CAZyme repertoire was used to elucidate the trophic lifestyle for *P. maydis*. The relative centroid distance (RCD) values are reported for each classification (on a scale from 0 to 1; Hane et al., 2020). An RCD value of 1 indicates the closest and most likely trophic classification while 0 indicates the farthest and least likely trophic classification (Hane et al., 2020).

|  | CAZymes | Predicted secreted CAZymes | CAZyme Class |  |  |  |  |  |
| --- | --- | --- | --- | --- | --- | --- | --- | --- |
|  |  |  | AA | CBM | GH | GT | CE | PL |
| *Phyllachora maydis* (PM02 assembly) | 336 | 92 | 54 | 20 | 153 | 78 | 31 | 0 |

**B**
| Trophic classification |  |  |  |  |
| --- | --- | --- | --- | --- |
| Saprotroph | Vasculartroph | Mesotroph | Monomertroph | Polymertroph |
| 0.88 | 0 | 0.42 | 1 | 0.43 |

### Selection of candidate effectors from *P. maydis* isolate PM02 (assembly ASM2933922v1)

Recent genomic and transcriptomic profiling of *P. maydis* revealed this fungal pathogen encodes 163 putative effector proteins, eighteen of which are abundantly expressed during disease development (Supplemental Table 1; MacCready et al., 2023). Given that these candidate effectors are upregulated during tar spot lesion development, we selected these effectors for further study. These eighteen candidate effectors ranged in size from 55 to 221 amino acids and three candidate effectors, PM02_g2959, PM02_g7163, and PM02_g7882, encode predicted nuclear localization signals (NLS) as determined by LOCALIZER (v1.0) (Table 2; Supplemental Table 1). Further examination of the protein sequences from two effector candidates, PM02_g378 and PM02_g8240, revealed that they encode a predicted GPI-anchoring domain and a superoxide dismutase domain, respectively (Table 2).

**Table 2.** *P. maydis* effector candidates investigated in this study.

| Protein ID<sup>a</sup> | Amino acids<sup>b</sup> | Protein size<sup>c</sup> | Predicted Localization<sup>d</sup> | Domain Annotation<sup>e</sup> |
| --- | --- | --- | --- | --- |
| PM02_g157 | 176 | 17.86 |  |  |
| PM02_g378 | 188 | 19.59 |  | GPI-anchoring (6-97) |
| PM02_g886 | 131 | 13.28 |  |  |
| PM02_1115 | 127 | 13.75 |  |  |
| PM02_1888 | 174 | 17.37 |  |  |
| PM02_g2094 | 94 | 10.39 |  |  |
| PM02_g2610 | 155 | 16.39 |  |  |
| PM02_g2959 | 62 | 6.59 | NLS (32-38) |  |
| PM02_g3916 | 121 | 12.74 |  |  |
| PM02_g4549 | 55 | 5.51 |  |  |
| PM02_g4592 | 129 | 13.97 |  |  |
| PM02_g4611 | 137 | 14.89 |  |  |
| PM02_g5788 | 138 | 14.14 |  |  |
| PM02_g7163 | 194 | 21.62 | NLS (67-81,134-149) |  |
| PM02_g7882 | 104 | 11.51 | NLS (94-102) |  |
| PM02_g7920 | 67 | 6.9 |  |  |
| PM02_g8240 | 221 | 23.18 |  | Superoxide dismutase-like (42-193) |
| PM02_g8507 | 61 | 7.07 |  |  |

a Protein IDs were mined from MacCready et al., (2023). Full-length protein sequences are indicated in Supplemental Table 1
b Numbers indicate the length of amino acids without the predicted signal peptide sequences
c Predicted molecular weight (kilodaltons; kDa) of the candidate effector proteins without the predicted signal peptide sequences
d Putative nuclear localization signals (NLS) were predicted by LOCALIZER (v1.0). Numbers within the parentheses correspond to amino acid positions
e Putative domains were identified using the InterPro database

### *In silico* structural modeling of *P. maydis* candidate effectors by AlphaFold2

Although we identified several candidate effectors with sequence-based domain predictions, effectors often lack conserved motifs as they rapidly evolve to evade host detection (de Guillen et al., 2015; Fouché et al., 2018). Recent work by Seong and Krasileva (2023) demonstrated that fungal pathogens often encode effectors with variable amino acid sequences but similar structures, wherein such effectors often share common protein folds. Hence, predicting the tertiary structure of fungal effectors can often provide insight into their putative functions (Seong and Krasileva, 2023). We, therefore, used AlphaFold2 to predict the tertiary structures of the candidate effectors (Jumper et al., 2021; Varadi et al., 2022). Visual inspection of the predicted protein structures revealed that several of our effector candidates had regions predicted with high confidence as determined by AlphaFold2 (Figure 1A). For example, the tertiary structure of the predicted copper/zinc superoxide dismutase-like (SOD) domain within PM02_8240 adopted a protein fold that was predicted with very high confidence (Figure 1A). Intriguingly, the amino-and carboxy-terminal regions within most of the candidate effectors were predicted with low confidence and primarily unstructured, which may suggest the presence of intrinsically disordered proteins regions (Figure 1A).

**Figure 1.**
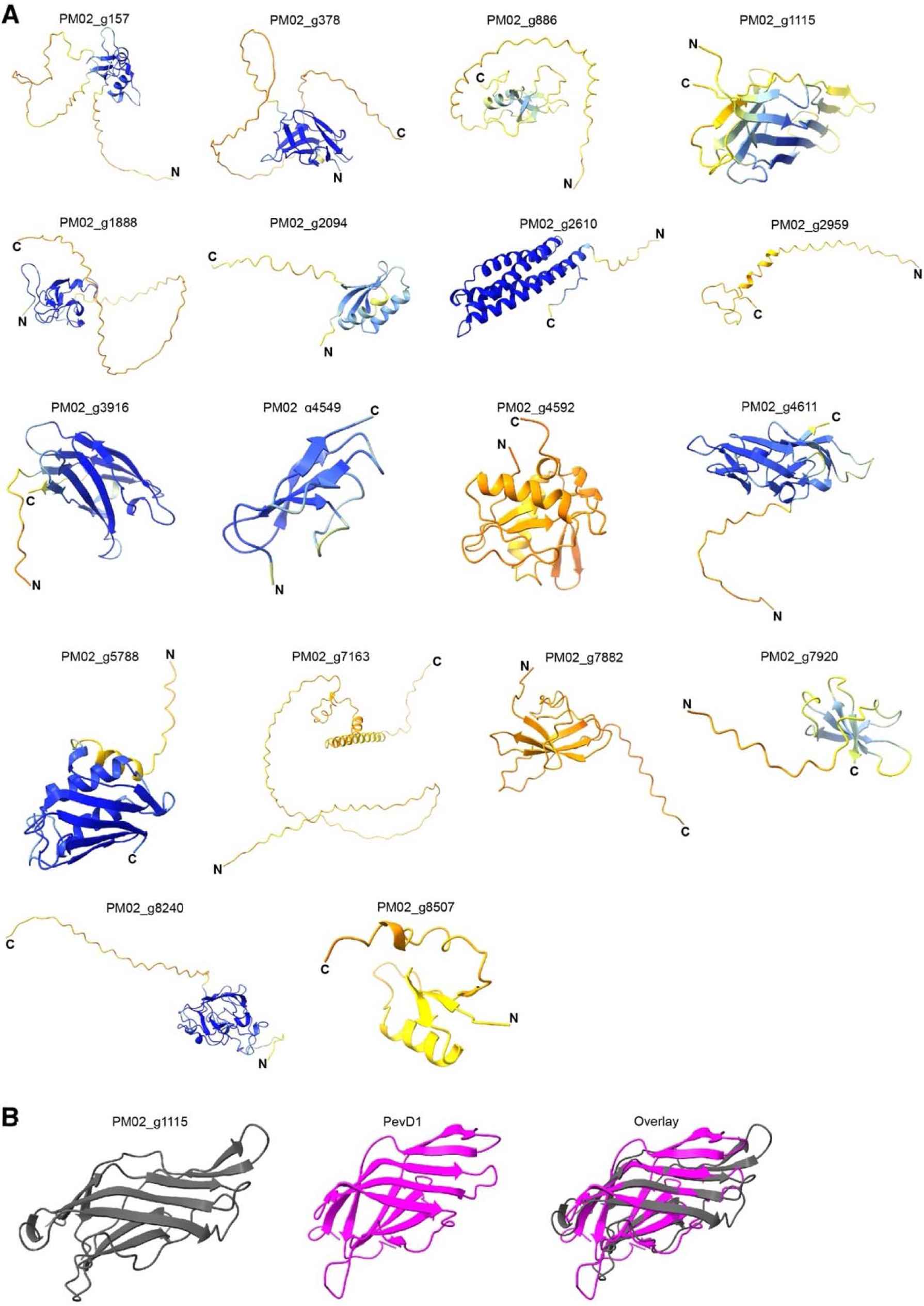
Predicted three-dimensional protein structures of the selected *P. maydis* candidate effectors as determined by ColabFold v1.5.2: AlphaFold2. A) Protein structures were predicted with ColabFold v1.5.2: AlphaFold2 and rendered using the ChimeraX-1.5 modeler package. Predicted protein structures were colored in ChimeraX-1.5 according to the AlphaFold2 pLDDT scores; predicted very high confidence regions are indicated by dark blue, high confidence regions are indicated by light blue, low confidence regions are indicated by yellow, and very low confidence regions are indicated by orange. N: amino terminus; C: carboxyl terminus. **B)** Foldseek analysis predicted structural similarity between PM02_g1115 (gray) and PevD1 (magenta) from *Verticillium dahlia* (PDB: 5XMZ). When overlaid (gray and magenta), these proteins appear to share six antiparallel β-sheets.

Having obtained the predicted protein structures of each of the effector candidates, we next utilized Foldseek to identify effectors with structural similarity to that of our subset of *P. maydis* candidate effectors (Supplemental Table 2; van Kempen et al., 2023). These analyses revealed that one candidate effector, PM02_g1115, adopted a predicted protein structure similar to that of the PevD1 effector from *Verticillium dahlia* (Figure 1B). PM02_g1115 encodes a β-barrel structure composed of ten β-strands as predicted by AlphaFold2 (Figure 1B). Structural alignment and examination of this *P. maydis* effector with PevD1 showed that PM02_g1115 shares six conserved anti-parallel β-sheets with PevD1 (Figure 1B). Further interrogation of the PevD1 protein showed that it is likely an *Alternaria alternata* 1 (Aa1)-like protein (Zhang et al., 2019). The first Aa1 protein was characterized from *Alternaria alternata* and described as having a β-barrel structure composed of 11 β-strands (Chruszcz et al., 2012). Aa1 proteins have been found exclusively in fungal species, specifically *Sordariomycetes* and *Dathideomycetes*, but their function in plant-fungal interactions have yet to be characterized (Chruszcz et al., 2012; Zhang et al., 2019).

### The majority of *P. maydis* candidate effector-fluorescent protein fusions localize to the nucleus and cytosol

Though our subset of putative effectors encode a signal peptide and are thus predicted to be secreted, it is unclear whether these proteins target specific subcellular compartments. We, therefore, generated candidate effector-super Yellow Fluorescent Protein (sYFP) fusions and assessed their subcellular localization patterns in *Nicotiana benthamiana* epidermal cells using laser-scanning confocal microscopy (Figure 2). Live-cell imaging revealed the majority of the *P. maydis* effector candidates evenly diffused throughout the nucleus and cytosol, similar to localization of the free sYFP control (Figure 3A). However, fluorescence signal from two candidate effectors, PM02_g378 and PM02_g2610, accumulated predominantly in the cytosol with weak fluorescence signal detected in the nucleus (Figure 3A).

**Figure 2.**
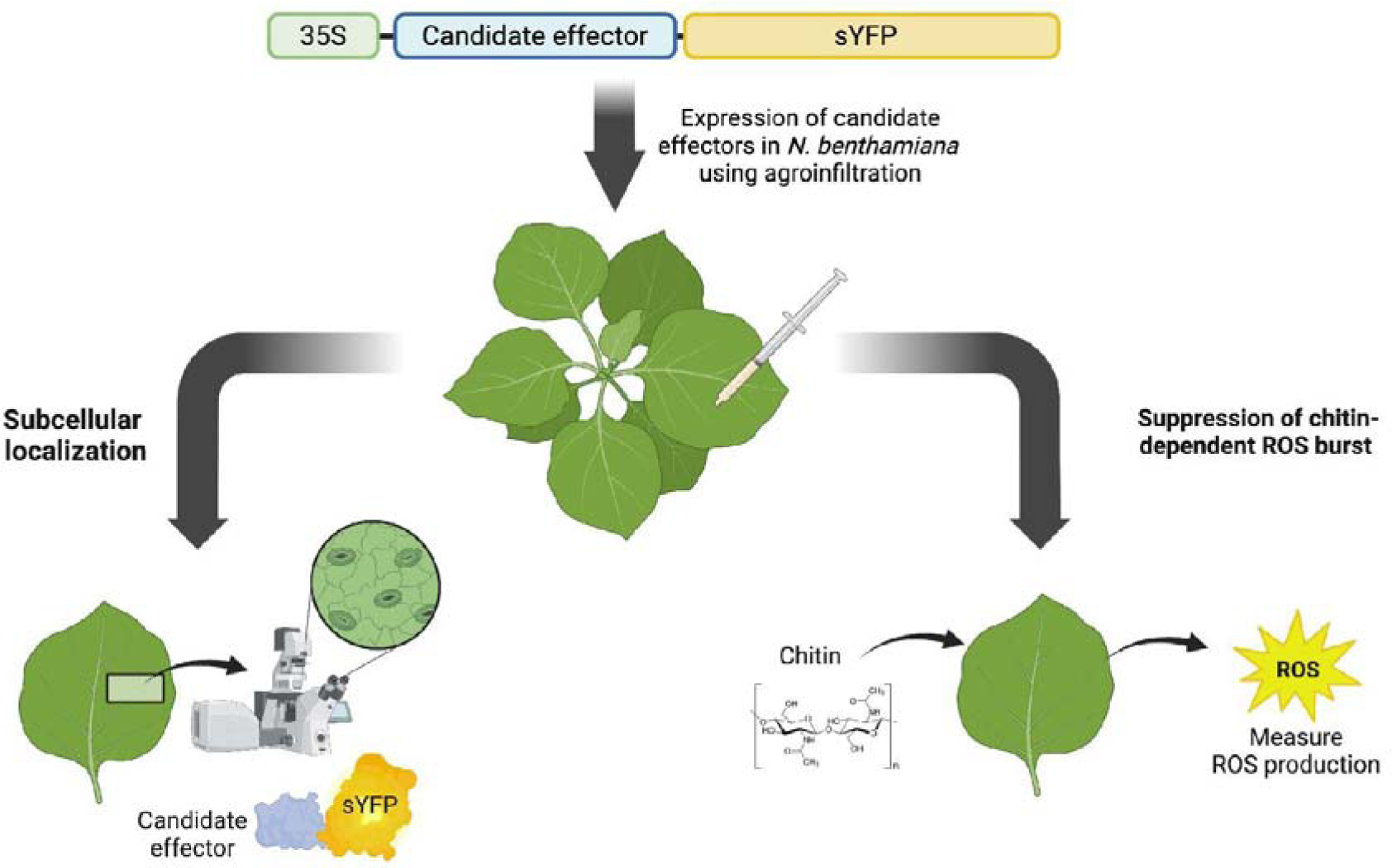
Schematic illustration of the experimental pipeline used to functionally characterize the selected *P. maydis* effector candidates in this study. The open reading frames of the selected *P. maydis* candidate effectors (without the signal peptide sequence) were fused to the N-terminus of super Yellow Fluorescent Protein (sYFP) and recombined into the pEG100 (pEarleyGate100) plasmid using Gateway cloning. The resulting constructs were transiently expressed in *Nicotiana benthamiana* to assess their subcellular localizations as well as to identify effector candidates that suppress chitin-mediated ROS production. Figure was created with Biorender (https://biorender.com).

**Figure 3.**
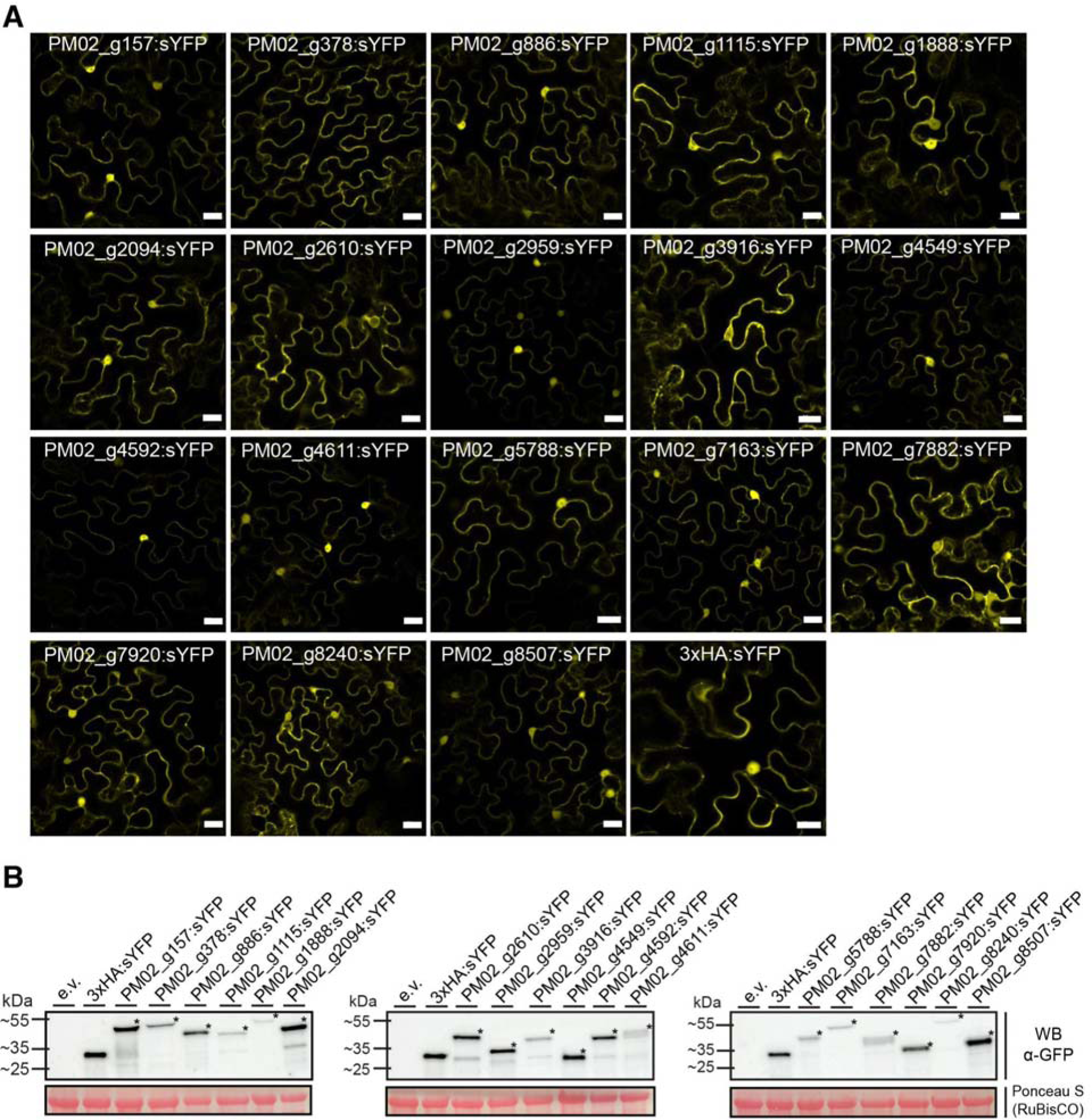
Subcellular localization and immunoblot analyses of the *P. maydis* candidate effectors in *N. benthamiana*. **A)** The indicated constructs were transiently expressed in *N. benthamiana* using agroinfiltration and *in planta* localization was assessed 24 hours post-agroinfiltration using laser-scanning confocal microscopy. Free sYFP (3xHA:sYFP) was used as a reference for nucleo-cytoplasmic distribution. Confocal micrographs are of single optical sections and the scale bar is 20µM. **B)** Candidate effector-fluorescent protein fusions accumulate protein in *N. benthamiana*. Total protein was extracted 24 hours following agroinfiltration in *N. benthamiana*. Equal volumes of total protein were resolved on 4-20% SDS-PAGE gels, blotted onto nitrocellulose membranes, and probed using anti-GFP antibodies. Free sYFP and empty vector (e.v) were included as controls. Ponceau staining of the RuBisCO large subunit was used as a loading control. Black asterisks denote the theoretical size of each fusion protein.

To confirm the integrity of the fusion proteins, we performed immunoblotting assays using anti-GFP antibodies (Figure 3B). Consistent with the subcellular localizations, all eighteen effector candidates accumulated detectable protein in *N. benthamiana* (Figure 3B). In addition to the band signal at the theoretical size of the protein fusions, some fusions showed a protein product at a lower molecular weight that coincided with free sYFP, suggesting cleavage of sYFP from the recombinant protein may have occurred (Figure 3B). Moreover, we observed that sYFP-tagged PM02_g4611 and PM02_g7882 consistently produced two distinct protein bands, suggesting these candidate effectors may undergo post-translational modifications (Figure 3B).

### A subset of *P. maydis* candidate effectors attenuate chitin-mediated ROS production

To gain initial insight into the putative functions of these eighteen effector candidates, we investigated whether any attenuate reactive oxygen species (ROS) production induced by chitin (Figure 2). To induce disease, fungal phytopathogens often rely, in part, on effectors that subvert early plant defense responses including the production of host-derived ROS (Chen et al., 2022; Navarrete et al., 2021; Saado et al., 2022). Given the functional importance of ROS accumulation in host immunity, we hypothesized that our *P. maydis* candidate effectors will also suppress PAMP-elicited ROS accumulation. To test our hypothesis, we transiently expressed each *P. maydis* candidate effector-fluorescent protein fusions in *N. benthamiana*. Plants were subsequently challenged with chitin and ROS production was monitored using a luminol-based assay. Consistent with our hypothesis, transient expression of three candidate effectors, PM02_g1115, PM02_g7882, and PM02_g8240, rapidly attenuated maximum chitin-mediated ROS production when compared to the free sYFP control in *N. benthamiana* (Figure 4A). To rule out the possibility that the lack of observed ROS suppression by some of the sYFP-tagged candidate effectors was due to insufficient protein accumulation, we performed anti-GFP immunoblot analyses. All effector-fluorescent protein fusions expressed detectable protein with no obvious protein degradation forty-eight hours post-agroinfiltration, demonstrating that the lack of ROS suppression by some of the candidate effectors is likely not a result of insufficient protein accumulation (Figure 4B). We, therefore, conclude that the *P. maydis* candidate effectors, PM02_g1115, PM02_g7882 and PM02_g8240, attenuate chitin-triggered ROS production when transiently expressed in *N. benthamiana*, and could potentially be required for pathogenicity.

**Figure 4.**
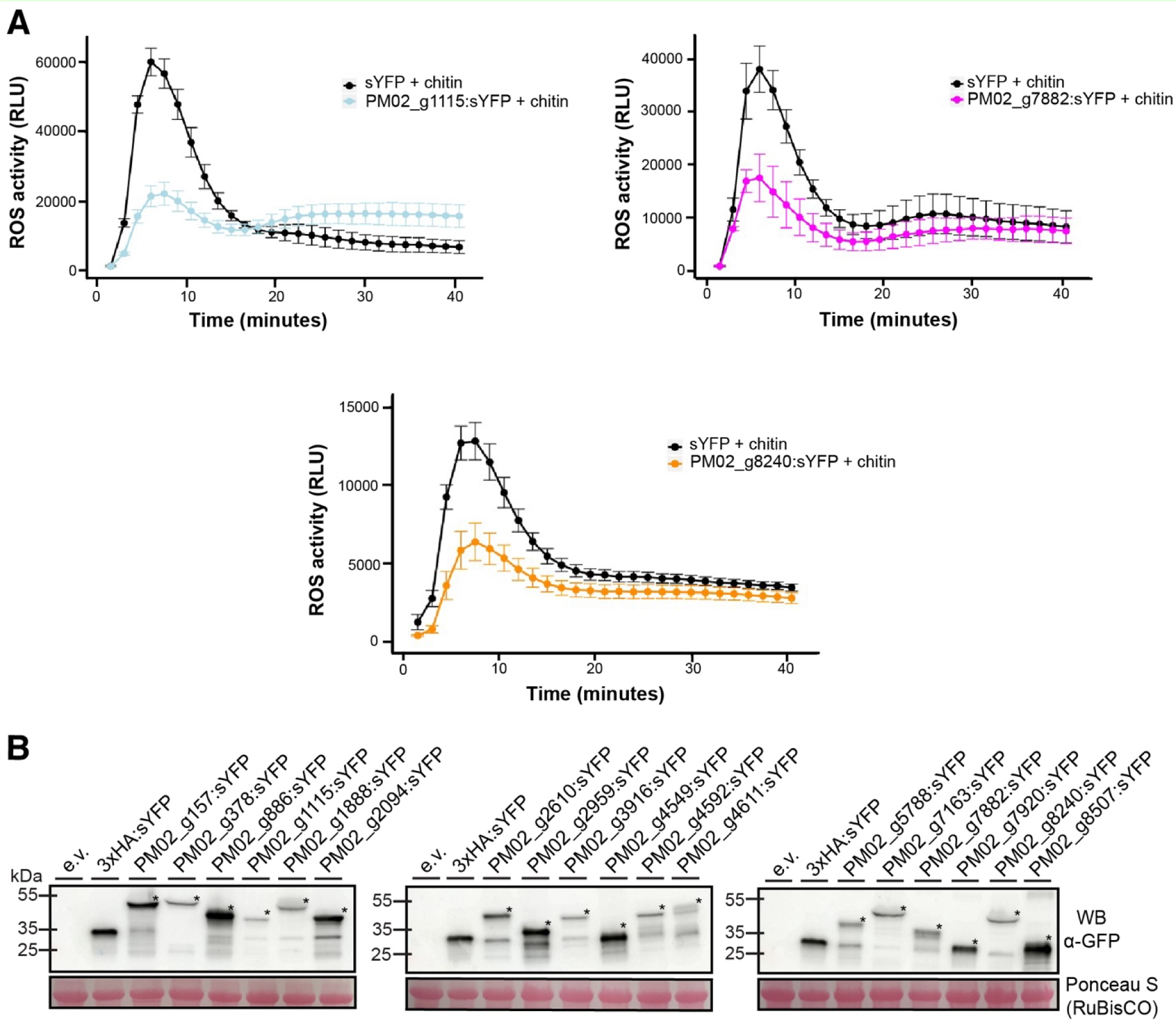
Examples of *P. maydis* candidate effectors that suppress chitin-dependent reactive oxygen species (ROS) burst in *N. benthamiana*. A) Transient expression of candidate effector-fluorescent protein fusions suppress reactive oxygen species (ROS) burst triggered by chitin in *N. benthamiana*. The indicated constructs were expressed in *N. benthamiana*. Forty-eight hours post-agroinfiltration, *N. benthamiana* leaf disks were collected, challenged with chitin, and relative luminescence was monitored for 40 minutes using a luminol-based assay. Free sYFP (3xHA:sYFP) was used as a reference control. **B)** Immunoblot assays depicting protein expression of the sYFP-tagged candidate effectors at 48 hours following agroinfiltration. Equal volumes of total protein were resolved on 4-20% SDS-PAGE gels, blotted onto nitrocellulose membranes, and probed using anti-GFP antibodies. Free sYFP and empty vector (e.v) were included as controls. Ponceau staining of the RuBisCO large subunit was used as a loading control. Black asterisks denote the theoretical size of each effector-fluorescent protein fusion.

## DISCUSSION

The results presented herein, combined with the previous work of MacCready and colleagues (2023), provide strong evidence that *P. maydis* is likely a monomertroph that utilizes effector proteins to suppress immune responses. To the best of our knowledge, our work is the first to use a genome-informed approach to elucidate the predicted trophic lifestyle as well as functionally characterize a subset of effectors from this fungal pathogen. Our results presented herein will likely assist in further understanding the infection strategies used by *P. maydis* as well as support the development of targeted disease management strategies against this fungal pathogen.

The observation that the CAZyme repertoire of fungal phytopathogens is often correlated with infection strategy afforded us the opportunity to infer a trophic lifestyle for *P. maydis* (Hane et al., 2020). Leveraging the most recent *P. maydis* genome annotation and the CATAStrophy prediction tool, we show *P. maydis* is enriched in CAZymes consistent with that of non-haustorial monomertrophs (biotrophs) (Table 1). Consistent with our analyses, parasitic fungal pathogens within the order *Phyllachorales* are characterized as obligate biotrophs (Mardones et al., 2017). Further examination of the predicted *P. maydis* CAZyme repertoire revealed this fungal pathogen contains a relatively similar quantity of predicted glycoside hydrolases compared to other biotrophic pathogens such as *Melampsora larici-populina* and *Puccinia graminis f. sp. tritici* further supporting our notion that *P. maydis* is likely a biotroph (Duplessis et al., 2011).

Moreover, recent work by Caldwell et al., (2023) revealed that, during infection of host cells, fungal hyphae develop into reproductive structures that form around but not within the host vasculature further supporting our observation this fungal pathogen is not vasculartroph (Table 1). Though the CATAStrophy pipeline is useful for predicting the lifestyle and trophic classification of filamentous plant pathogens, this approach can provide low accuracy if the poor-quality genome assembly and inaccurate CAZyme predictions are used as inputs (Hane et al., 2020). Nevertheless, the genome assembly of isolate PM02 used for this pipeline from MacCready et al. (2023) has a BUSCO score of 98.6%, demonstrating a high-quality assembly of *P. maydis*.

The observation that fungal pathogens may encode sequence-unrelated structurally similar effectors encouraged us to predict the tertiary structures of the candidate effectors and perform structural comparative analyses. Here, we predicted the protein structures of each of our candidate effectors and identified a putative structural homolog of the PM02_g1115 effector using the structural alignment tool Foldseek (Figure 1). Intriguingly, the Foldseek analyses revealed the *P. maydis* effector PM02_g1115 may be structurally analogous to the PevD1 effector from *V. dahlia*. PevD1 is a secreted Alt a 1 (Aa1)-like protein that interacts with and subverts the antifungal activity of the pathogenesis-related protein GhPR5 from cotton (*Gossypium hirsutum*), thereby facilitating *V. dahliae* disease development (Zhang et al., 2019). Importantly, knockout of PevD1 expression reduces *V. dahliae* virulence in cotton, demonstrating PevD1 is likely required for wilt symptom development (Zhang et al., 2019). Additionally, application of purified PevD1 protein to cotton seedlings prior to *V. dahliae* inoculation improved resistance against symptom development (Bu et al., 2014). Though PevD1 is the only characterized Aa1-like protein from *V. dahliae*, Aa1-like proteins are present in other fungal phytopathogens including *Magnaporthe oryzae* (Zhang et al., 2019). For example, MoHrip1 is an Aa1-like protein from *M. oryzae* that localizes to the host cell periphery and activates several immune responses including ROS production and HR-like cell death, revealing MoHrip1 likely functions as a PAMP (Zhang et al., 2017). Taken together, these observations suggest fungal phytopathogens, including *P. maydis*, encode Aa1-like proteins that modulate host immune responses. Future research will thus aim to identify the host targets of PM02_g1115 and seek to address the functional role of this effector in *P. maydis* disease development.

Though our study relied on heterologous overexpression of epitope-tagged effector proteins, the observation that several candidate effectors consistently attenuated chitin-mediated ROS production when expressed within the plant cell suggests these proteins are likely secreted, *bona fide* effectors (Figure 4). Consistent with our observations, Navarrete et al., (2021) showed that the *U. maydis* effectors Tay1 and Mer1 inhibit PAMP-triggered ROS burst when transiently expressed in *N. benthamiana* and maize. However, fusing a nuclear localization signal to Tay1 and a nuclear export signal to Mer1, respectively, abolished their ROS inhibiting activities (Navarrete et al., 2021). Additionally, the *U. maydis* effector Rip1 suppresses flagellin22 (flg22)-triggered ROS burst, and such immune suppressing activities was independent of Rip1 *in planta* subcellular localization (Saado et al., 2022). Hence, investigating the subcellular localization patterns of fungal effectors in parallel with testing for immune suppression activities provides foundational insight into host proteins potentially targeted by fungal effectors. Future work will thus address whether the immune suppression activities of PM02_g1115, PM02_g7882 and PM02_g8240 is dependent upon their subcellular localizations as well as investigate if these candidate effectors suppress additional immune responses. To this end, the type III secretion system from the bacterial pathogen *Pseudomonas syringae* pv. *tomato* DC3000 has been previously used as a surrogate to deliver fungal effectors into plant cells to ascertain effector functionality (Dalio et al., 2018; Fabro et al., 2011; Sohn et al., 2007). For example, the wheat stripe rust effector Shr7 was shown to suppress the cell death responses elicited by *P. syringae* pv. *tomato* DC3000 in wheat (Ramachandran et al., 2017; Yin and Hulbert, 2010). Furthermore, Qi and colleagues (2016) identified an effector protein from *Phakopsora pachyrhizi* termed PpEC23 that suppresses *P. syringae* pv. *tomato* DC3000-mediated immune responses in soybean. These prior studies thus demonstrate that *P. syringae* pv. *tomato* DC3000 is an effective heterologous expression system for delivering non-bacterial effectors into host cells (Fabro et al., 2011; Sohn et al., 2007). Recent work by Jaiswal and Helm (2023) showed *P. syringae* pv. *tomato* DC3000 induces defense responses in diverse maize inbreds, including hydrogen peroxide accumulation and an increase in transcript expression of maize defense genes. Hence, future studies should aim to use *P. syringae* pv. *tomato* DC3000 as a surrogate for heterologous expression to assess whether any of the candidate effectors suppress *P. syringae* pv. *tomato* DC3000-mediated immune responses in maize. Alternatively, generation of transgenic maize overexpressing epitope-tagged candidate effectors would also provide a direct test of their role in pathogenicity.

Though *P. maydis* is considered one of the most important foliar pathogens of maize, our knowledge of the trophic lifestyle and functional role of effector proteins from this fungal pathogen remains limited. In our study, we combined state-of-the-art computational methods, advanced protein structure prediction algorithms, and experimental approaches to further elucidate the trophic lifestyle and unveiled putative functions of effector proteins from *P. maydis*. We predict our work will be instrumental in guiding additional research aimed at elucidating the potential molecular and cellular mechanisms of action used by this fungal pathogen to induce tar spot in maize.

## ACKNOWLEDGEMENTS

We thank Dr. Tesfaye Mengiste (Purdue University) for access to the microplate reader for the ROS suppression assays. We would also like to thank Dr. Terri Cameron, Brendan Lane, and Dr. Katherine Rivera-Zuluaga for technical assistance, and the Purdue University Imaging Facility for access to the Zeiss LSM880 Axio Examiner upright confocal microscope. The authors would also like to thank Dr. Ariel Sturgill, Dr. Rajdeep Khangura, and Dr. Jose Salguero-Linares for insightful discussions and critical reading of the manuscript.

## SUPPLEMENTARY TABLES

**Supplemental Table 1:**
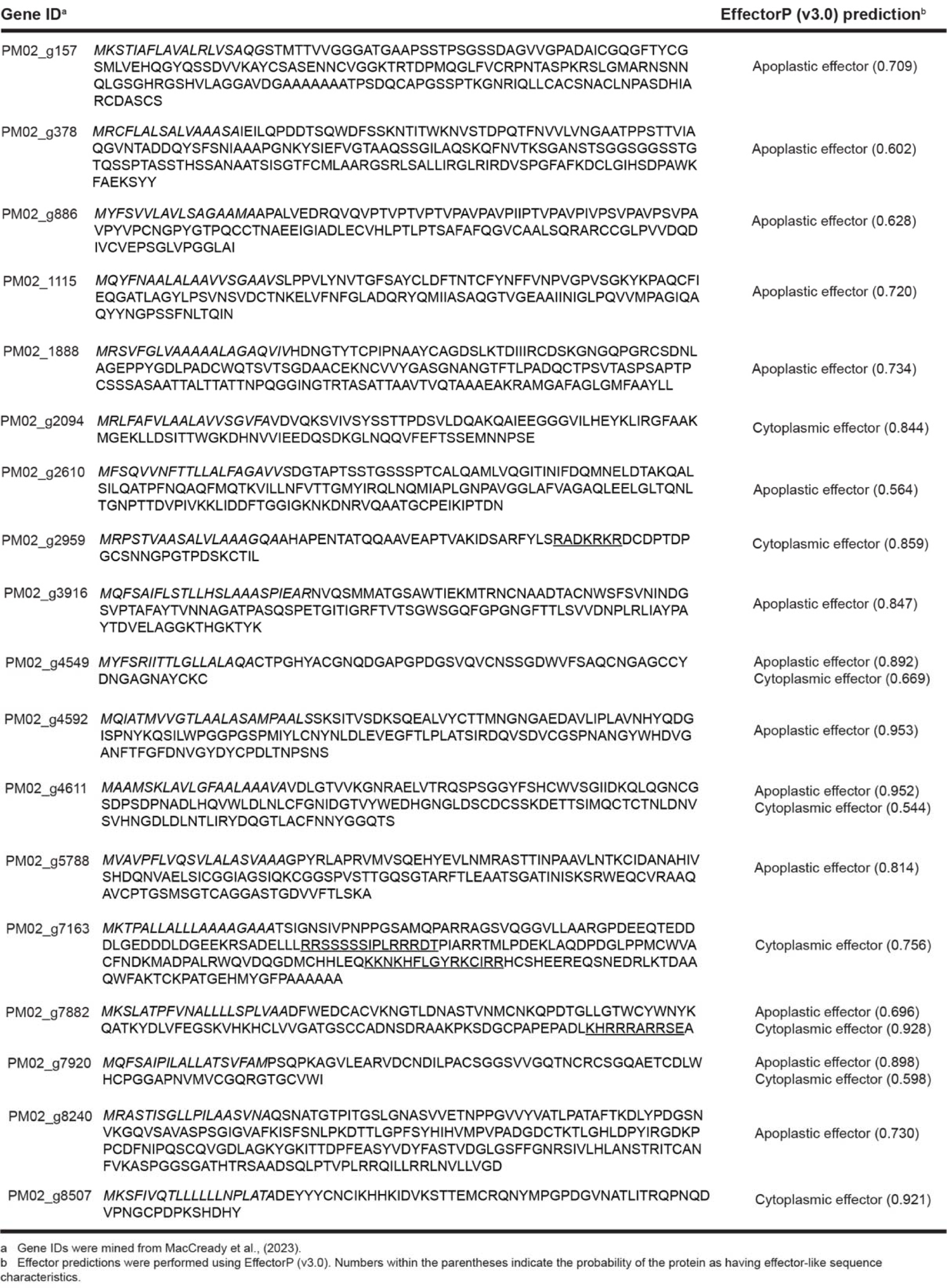
Protein sequences of the 18 *P. maydis* candidate effectors. Italicized and underlined amino acids represent the predicted signal peptide sequences and putative nuclear localization signals, respectively.

**Supplemental Table 2:**
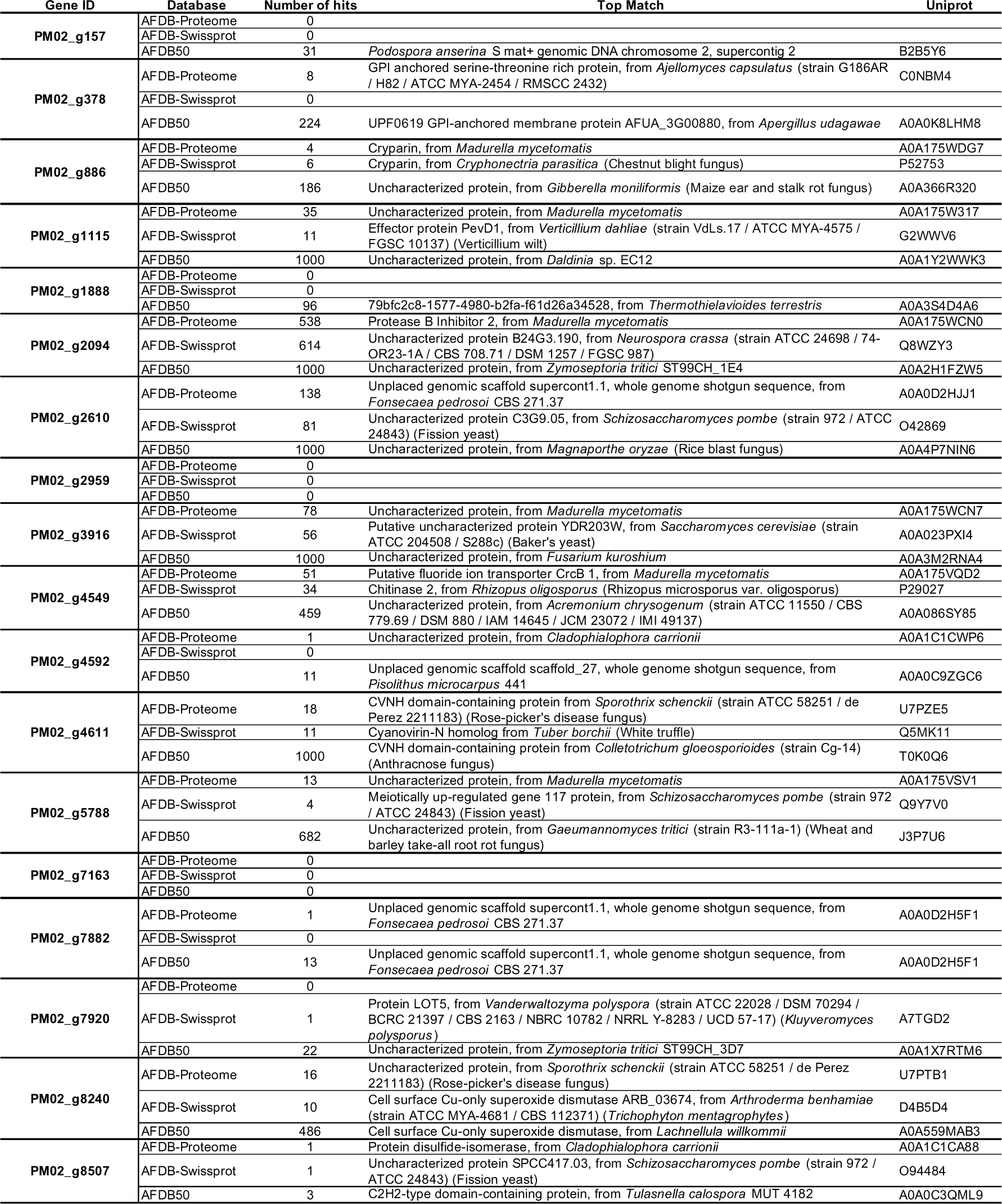
Results from the Foldseek analyses. Predicted structural matches are presented for each of the *P. maydis* candidate effectors investigated in the current study. Predicted matches are presented from each iteration of the AlphaFold Protein Structure Database (AFDB-Proteome, AFDB-SwissProt, AFDB50) and their corresponding UniProt ID is provided.

